# Dynamic balance of H3K9me2 heterochromatin by CoREST-2 and RE-1 in growing neurons

**DOI:** 10.1101/2024.12.31.629745

**Authors:** José F. Drube, Andrés Cardozo-Gizzi, Victoria Rozés-Salvador, Imanol Martínez, Danilo G. Ceschin, Laureano Giordano, Lucas Serniotti, Laura Gastaldi, Mónica Remedi, Ana L. Moyano, Alfredo Cáceres, Carlos Wilson

## Abstract

A well-balanced chromatin dynamic is vital for the survival and physiology of all cells, including brain neurons. Heterochromatin, commonly associated with transcriptional silencing, also plays significant roles in maintaining genomic stability and facilitating DNA-repair processes. Notably, the bimethylation of histone H3 at lysine 9 (H3K9me2), a hallmark of repressive heterochromatin, supports the axonal specification of neurons. However, neuronal maintenance of H3K9me2 equilibrium remains understudied.

In this work, we unveil a dynamic equilibrium of H3K9me2 regulated by the epigenetic factor CoREST-2 and RE-1 DNA motifs, sustaining axonal and dendritic outgrowth. Using primary cultures of rat hippocampal neurons and a combination of advanced imaging techniques, we observed an enriched nuclear accumulation of CoREST-2 and H3K9me2 along neuronal development. Genetic silencing of CoREST-2 induced axon-dendrite retraction, accompanied by an increase in nuclear levels of H3K9me2. To further investigate heterochromatin structure at the nanoscale, we employed STED nanoscopy and discovered that H3K9me2 is organized into small nanodomains, which were notably enlarged following the suppression of CoREST-2. In contrast, the genetic blockade of RE-1 DNA motifs led to axon-dendrite retraction alongside the disassembly of H3K9me2 nanodomains.

These findings highlight that CoREST-2 and RE-1 sites actively shape neuronal H3K9me2 heterochromatin. Moreover, they uncover that maintaining a precise balance of H3K9me2 is essential for the extension of axons and dendrites, underpinning the connectivity and plasticity of brain neurons.

## Introduction

Chromatin regulation, and particularly histone post-transcriptional modifications (PTM), is crucial for the life of all eukaryotic cell types. In the nervous system, histone PTM contributes to neurogenesis, neuronal polarization, axon pathfinding, and regeneration ^1–7^. In particular, the bimethylation of lysine 9 in histone 3 (H3K9me2) is a repressive signature at the heterochromatin enabling post-mitotic development of brain neurons ^8,9^.

The homeostasis of H3K9me2 depends on the concerted action of histone methyltransferases and lysine-specific demethylases, such as G9a and LSD1, respectively ^8–11^. Both are recruited to the chromatin by additional factors, including the family of CoREST proteins or the transcriptional repressor REST that acts through a DNA motif called Regulatory Element-1 (RE-1) ^10,12,13^. A significant part of this evidence has been obtained in non-neuronal cells. Consequently, their contribution to fine-tune H3K9me2 heterochromatin in brain neurons has remained understudied.

The family of CoREST proteins involves 3 paralogues ^14^. CoREST-1, the most studied member of the family, is expressed in embryonic and post-mitotic mouse brains, and its suppression halts the migration of neurons from the ventricular zone to the cortical plate of the brain cortex ^15^. Immunoprecipitation of CoREST-1 followed by enzymatic assays suggested histone-methyltransferase activity, most likely by G9a recruitment ^12,16^. By contrast, neuronal functions of CoREST-2 and -3 are unknown. In embryonic stem cells and neural precursors, CoREST-2 binds to the lysine specific demethylase 1 (LSD-1), an enzyme linked to H3K4 demethylation ^17^. Whether CoREST-2 modulates H3K9me2 homeostasis in brain neurons remains to be determined.

Regulation of H3K9me2 heterochromatin is also linked to the transcriptional repressor REST/NRSF and DNA motifs called Regulatory Elements 1 (RE-1, also NRSE) ^16,19^. In rodents, REST binds to RE-1 sites, a structural platform of 20-25 nucleotides used for epigenetic silencing ^20,21^. While REST is highly expressed in stem cells to prevent neuronal differentiation by repressing genes harboring RE-1 sites, its abundance drops to low-to-undetectable levels in post-mitotic neurons. Consequently, neuron-specific genes are unrepressed, enabling neuronal differentiation ^22^. RE-1 sites were initially described in neuronal genes, such as BDNF, Nav1.2, SGC10, and synaptotagmin-4, among others ^20,21,23^. However, i*n silico* analyses support the notion that more than 1,000 genes of the mouse and human genome harbor RE-1 sites, and their functions are not strictly related to neuron-specific roles ^24^. Whether RE-1 sites remain active in post-mitotic neurons is a matter of debate, and a potential contribution to chromatin homeostasis and neuronal growth needs to be determined.

This work unveils two new mechanisms controlling H3K9me2 homeostasis. Using primary cultures of rat hippocampal neurons along with super resolution microscopy, we report that nuclear depletion of CoREST-2 enlarges H3K9me2 heterochromatin nanodomains and restrains axon-dendrite growth. Additionally, genetic blockade of RE- 1 sites, by expressing the mutant REST-DBD, dismantles H3K9me2 heterochromatin nanodomains, paralleled by axon and dendrite retraction. Together, these findings suggest that CoREST-2 and RE-1 coordinate mechanisms maintaining H3K9me2 heterochromatin homeostasis in post-mitotic neurons. Moreover, it supports the idea that a precise balance of H3K9me2 is required to preserve axon-dendrite extension, which is crucial for maintaining the connectivity of brain neurons.

## Results

### Nuclear accumulation of CoREST-2 safeguards neuronal growth

Genetic suppression of CoREST-2 in mouse brains, either using knock-out mice or *in utero electroporation* knock-down assays, reported a reduction on the number of neuronal progenitors and neurons ^17,25^. Still, its contribution to H3K9me2 heterochromatin homeostasis and nerve cell growth in post-mitotic neurons remains unknown.

To this end, the expression and nuclear destination of CoREST-2 was evaluated by immunofluorescence (IF) staining and confocal z-stack imaging in embryonic rat hippocampal neurons cultured for 1, 3, 7 and 14 days in vitro (DIV) (Figure 1A, D-F). At 14 DIV, neurons also exhibited nuclear accumulation, but an increase in the fluorescence signal was also detected outside the cell nucleus. Consequently, the nuclear/cytosolic ratio peaked by 7 DIV to then decay at 14 DIV, mostly due to the cytosolic pool of CoREST-2 at this timepoint (Figure 1F).

**Figure 1.**
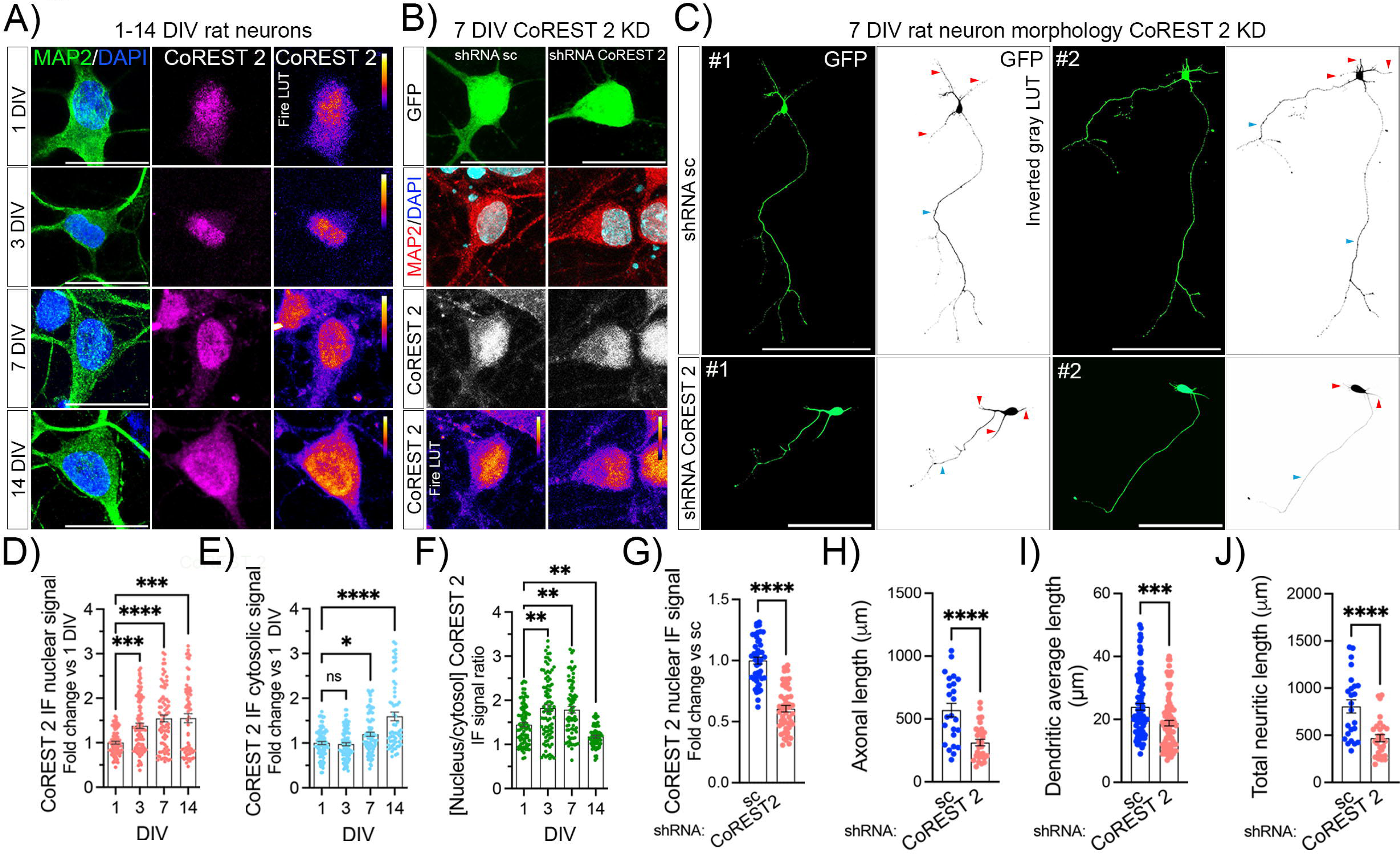
Nuclear accumulation of CoREST-2 safeguards neuritic growth. **A)** Representative images of rat hippocampal neurons cultured for 1, 3, 7 and 14 DIV, and stained for MAP2 (neuronal marker, green), CoREST-2 (magenta) and DAPI (nuclear region, blue). Fire LUT for CoREST-2 is shown to visualize fluorescence intensity. Images represent confocal z-stack maximal intensity projection. Scale bar: 10 µm. **B)** Representative images showing control and CoREST-2 KD neurons. Neurons were transfected with shRNA sc or shRNA CoREST-2 plasmids 18 h after plating and fixed at 7 DIV. Plasmids encode GFP for visualization of transfected cells. Staining: MAP2 (red), DAPI (blue) and CoREST-2 (gray and Fire LUT). Images represent confocal z-stack maximal intensity projection. Scale bar: 10 µm. **C)** Morphologies of 7 DIV control and CoREST-2 KD neurons. Images show z-stack maximal fluorescence intensity of GFP. Two representative cases are shown by condition. Scale bar: shRNA sc: 200 µm; shRNA CoREST-2: 100 µm. **D)** Quantification of nuclear CoREST-2 IF nuclear intensity. Kruskal- Wallis test, Dunn’s multiple comparison post-test, N=3. **E)** Quantification of cytosolic CoREST-2 IF intensity. Kruskal-Wallis test, Dunn’s multiple comparison post-test, N=3. **F)** CoREST-2 nuclear enrichment relative to cytosol IF. Kruskal-Wallis test, Dunn’s multiple comparison post-test, N=3. **G)** Estimation of CoREST-2 KD efficiency by quantifying nuclear IF signal of CoREST-2 vs shRNA sc. Mann-Whitney test, N=3. **H-J)** Quantification of axon (H, blue arrowheads), dendritic (I, red arrowheads) and total neuritic length (J). Mann-Whitney test, N=3.

To evaluate the contribution of CoREST-2 to neuronal growth, 1 DIV neurons were transfected with a plasmid encoding a small hairpin RNA (shRNA) targeting CoREST-2 mRNA and fixed at 7 DIV (Figure 1B, G). The green fluorescent protein (GFP) encoded in the pCAGIG backbone was used as a reporter ^8,26^. Considering the nuclear enrichment of CoREST-2 at this timepoint, the knock-down (KD) efficiency was estimated by a whole-nucleus z-stack imaging (Figure 1B, G). Accordingly, CoREST-2 KD neurons displayed 40% reduction of the nuclear IF signal (Figure 1G). Moreover, the phenotype of CoREST-2 KD neurons severely affected process extension, reducing both axonal and dendritic lengths by 50% and 30%, respectively (Figure 1C, H-J). Thus, this data indicates that CoREST-2 is crucially required for the growth of post mitotic rat neurons.

### CoREST-2 and H3K9me2 share common chromatin nanodomains

Considering the evidence linking CoREST proteins, G9a and H3K9 methylation, we wondered about a hypothetical crosstalk to maintain H3K9me2 homeostasis. Nuclear levels of G9a and H3K9me2 were determined at 1,3,7 and 14 DIV neurons by IF staining and confocal z-stack imaging (Figure 2A-F). Neurons exhibited a progressive enrichment of G9a and H3K9me2, reaching maximal values after 7 DIV. Therefore, this data reinforces the idea that CoREST-2, G9a, and H3K9me2 accumulate in neuronal nuclei within the first two weeks of development.

**Figure 2.**
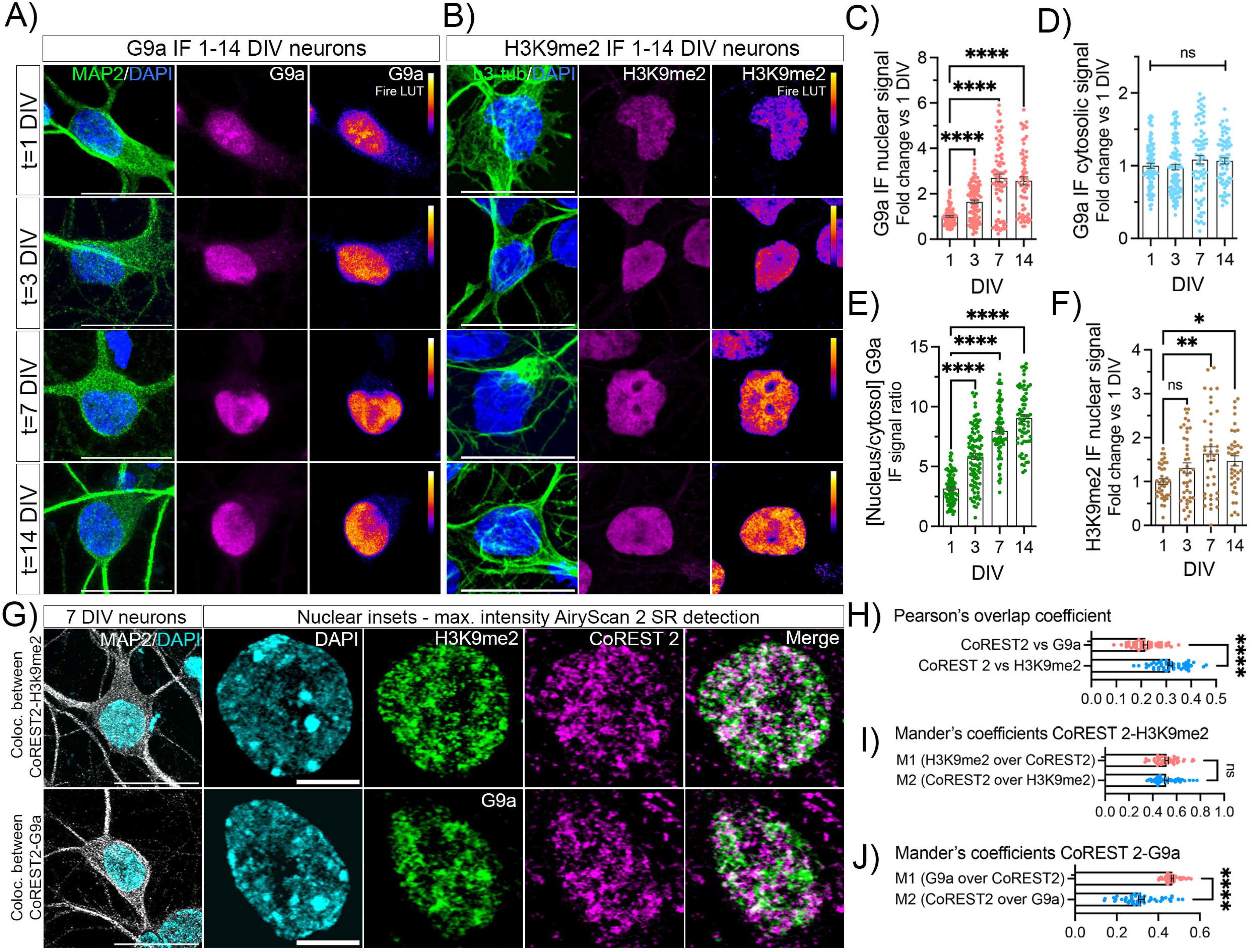
Spatial correlation between CoREST-2, H3K9me2 and G9a. **A, B)** Representative images of rat hippocampal neurons cultured for 1, 3, 7 and 14 DIV, and stained for MAP2 (A, neuronal marker, green), b3-tubulin (B, neuronal marker), G9a or H3K9me2 (A, B, respectively; magenta) and DAPI (nuclear region, blue). Fire LUT for G9a and H3K9me2 is shown to visualize fluorescence intensity. Images represent confocal z-stack maximal intensity. Scale bar: 10 µm. **C)** Quantification of nuclear G9a IF nuclear intensity. Kruskal-Wallis test, Dunn’s multiple comparison post-test, N=3. **D)** Quantification of cytosolic G9a IF nuclear intensity. Kruskal-Wallis test, Dunn’s multiple comparison post-test, N=3. **E)** G9a nuclear enrichment relative to cytosol. Kruskal- Wallis test, Dunn’s multiple comparison post-test, N=3. **F)** Quantification of nuclear H3K9me2 IF nuclear intensity. Kruskal-Wallis test, Dunn’s multiple comparison post- test, N=3. **G)** Representative images of 7 DIV neurons co-stained either with CoREST-2 and H3K9me2 or CoREST-2 and G9a. Images represent z-stack maximal intensity using LSM 980 AiryScan 2 microscope. Staining: MAP2 (gray), DAPI (blue), H3K9me2 or G9a (green), CoREST-2 (magenta). Scale bar: neuronal body: 10 µm; nuclear insets: 5 µm. **H)** Pearson’s overlap coefficient between CoREST-2, H3K9me2 and G9a. Mann-Whitney test, N=3. **I)** Mander’s colocalization coefficients for the pair CoREST-2 – H3K9me2. Mann-Whitney test, N=3. **J)** Mander’s colocalization coefficients for the pair CoREST-2 – G9a. Mann-Whitney test, N=3.

Using Airyscan microscopy, an advanced confocal technique that improves resolution and signal-to-noise ratio, we found that CoREST-2 exhibits a punctate pattern in 7 DIV neuronal nuclei (Figure 2G, upper and lower panel). Moreover, a similar non-homogenous distribution was observed for H3K9me2 and G9a (Figure 2G), unveiling differences compared to regular confocal imaging (Figure 2A-B). These observations are consistent with previous works using different cell lines and mouse brains ^27^.

CoREST-2 overlapped with H3K9me2 in the nucleus of 7 DIV rat neurons, visualized by white spots in the merge image (Figure 2G, upper panel). This association represented almost 30% of the area analyzed, according to Pearson’s coefficient (Figure 2H). Additionally, Mander’s coefficients for the pair CoREST-2-H3K9me2 suggested 40% of overlap between channels (Figure 2I). In contrast, Pearson’s coefficient for the pair CoREST-2 – G9a reported 20% overlap (Figure 2G, H), and disparities between M1 and M2 coefficients were found. Notably, M2 indicates that only a small fraction of CoREST-2 domains co-distributes with G9a (Figure 2J). Collectively, this data suggests that CoREST-2 associates to H3K9me2 heterochromatin in rat neurons.

### CoREST-2 preserves H3K9me2 heterochromatin nanodomains

Considering the association between CoREST-2 and H3K9me2 heterochromatin, we explored the possibility of a functional link between them. Accordingly, 1 DIV neurons were transfected with the plasmid shRNA CoREST-2 and fixed at 7 DIV, followed by IF staining and nuclear z-stack imaging (Supplementary Figure 1). As a result, CoREST-2 KD neurons exhibited a gain of approximately 20% of H3K9me2 IF nuclear signal, suggesting that physiological functions of CoREST-2 correlate with histone demethylase activity.

To further analyze H3K9me2 at heterochromatin organization, neuronal nuclei were visualized using stimulated emission depletion (STED) microscopy, improving spatial resolution down to the nanoscale ^28,29^. Figure 3A shows the nuclei of 7 DIV control and CoREST-2 KD neurons, transfected as indicated. This analysis provided us with information at several layers. For instance, the segmentation analysis of STED images revealed that H3K9me2 is organized in small nanodomains averaging a median equivalent area diameter of 60 nm (Figure 3A, C). Control and CoREST-2 KD neurons displayed a mean of 2,000 domains per nucleus, without changes between conditions. Nevertheless, CoREST-2 KD neurons displayed a significant increase of IF intensity in H3K9me2 nanodomains (Figure 3B), which is consistent with the whole-nucleus z-stack imaging addressed by confocal microscopy shown in the Supplementary Figure 1.

**Figure 3.**
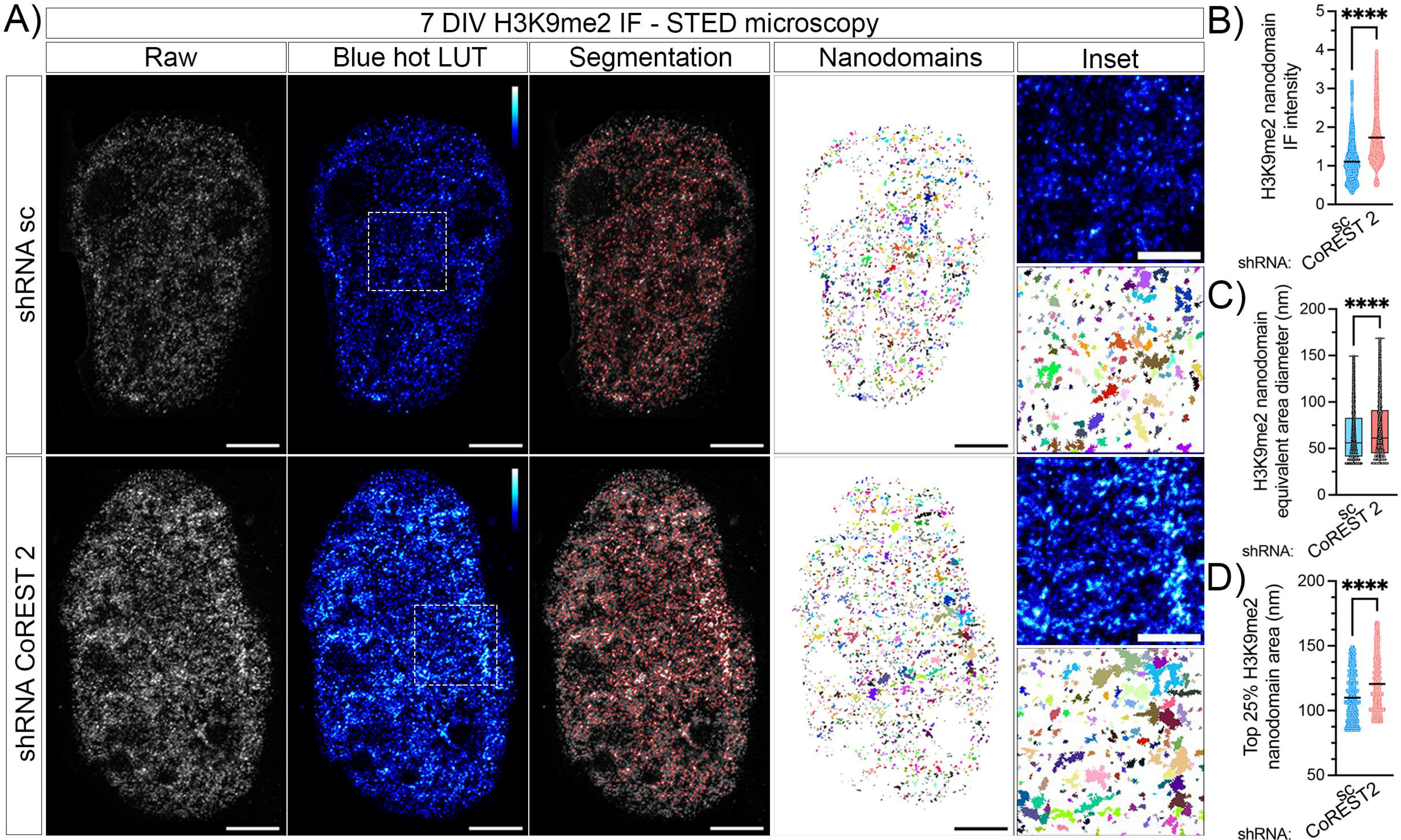
Silencing CoREST-2 enlarges H3K9me2 nanodomains in rat neurons. **A)** H3K9me2 nuclear detection in control and CoREST-2 KD neurons by STED nanoscopy. Neurons were transfected 18 h after plating, fixed at 7 DIV and stained to detect H3K9me2 by STED nanoscopy. The blue hot LUT is shown to visualize IF intensity. “Segmentation” shows the endpoint of the pipeline for domain segmentation, whilst “Nanodomains” the result. Colors do not mean intensity, but a visual representation of H3K99me2 domains. Scale bar: 2 µm; insets: 1 µm. **B)** IF intensity measured in each H3K9me2 nanodomain of control and CoREST-2 KD neurons. Mann-Whitney test, N=3. **C)** H3K9me2 equivalent area diameter of control and CoREST-2 KD neurons (whole population). Kolmogorov-Smirnov test, N=3. **D)** Representation of the top 25% largest H3K9me2 nanodomains of control and CoREST-2 KD neurons (equivalent area diameter). Kolmogorov-Smirnov test, N=3.

Of note, quantification of IF signal in STED nanodomains reported a higher increase compared to confocal whole-nucleus analysis, suggesting local changes in H3K9me2 heterochromatin after silencing CoREST-2 (Figure 3B vs Supplementary Figure 1B).

Additionally, the area of each segmented nanodomain was also measured (Figure 3C). For this parameter, we observed a slight but significant increase on the equivalent area diameter in CoREST-2 KD neurons. The blox plot shown in Figure 3C represents the global distribution of control and CoREST-2 KD neurons, suggesting an enlargement of the nanodomains belonging to the highest quartile in CoREST-2 KD neurons (top 25%). To clearly dissect this population, the Figure 3D plots the distribution of the top 25% of each sample, reinforcing that CoREST-2 KD neurons displayed an increase of the largest H3K9me2 nanodomains.

### RE-1 sites preserve neuronal growth and H3K9me2 nanodomains organization

Considering the experimental approach used in this work, we decided to revisit the functional role of regulatory elements 1 (RE-1) in maintaining chromatin homeostasis in neurons. RE-1 are consensus DNA sites recognized by the transcriptional repressor REST/NRSF ^20,21,23^. During neurogenesis, REST is quickly repressed to enable the differentiation of neural precursors into neurons ^22^. In 2004, an *in silico* and biochemical approach reported that more than 1,000 genes of the human and mouse genome harbor RE-1 sites ^24^. Following this track, we addressed a gene ontology (GO) analysis based on an *in-silico* search of RE-1 sites in the rat genome (Supplementary Figure 2, SF2). The SF2A shows the workflow of the analysis, identifying 637 rat genes containing RE-1 sites, located within -1,500 base pairs (bp) of the transcription start site (TSS) with q- values < 0.05. We selected genes based on this criterion to obtain the most-likely RE-1 – dependent genes, although this number can be higher in case of less-stringent criteria. The SF2B shows the GO analysis, indicating molecular functions (MF) and biological processes (BP). As expected, BP are related to anatomical structure development, signaling and nervous system, aligning with the developmental role of RE-1 sites. However, GO analysis uncovered global BP such as vesicle-mediated transport, DNA-templated transcription, cytoskeleton organization and programmed cell death. The raw data of this analysis is available as a supplementary spreadsheet for case-by-case perusals, including RE-1 in the rat, mouse and human genome.

Still, whether RE-1 sites control H3K9me2 homeostasis is unknown. Therefore, we tested this hypothesis by expressing the REST DNA-binding domain (DBD) construct ^20,24,30,31^. This plasmid encodes a mutant form of REST harboring deletions at the N- and C-terminal domains, used for epigenetic repression, but preserving 8 of 9 zinc-finger domains needed for DNA binding. Consequently, the REST-DBD leads to epigenetic deregulation by competitive inhibition and blockade of RE-1 sites ^20,22,31^(20, 32).Accordingly, 1 DIV neurons were transfected with the REST-DBD and IF stained at 7 DIV for REST detection by z-stack confocal imaging (Supplementary Figure 3). A GFP-encoding plasmid was used as a reporter. Control neurons exhibited a faint nuclear signal for endogenous REST, which became evident after expressing the REST-DBD construct (Supplementary Figure 3A, B). In this regard, our data aligns with a well-documented body of evidence reporting that REST is strongly repressed in post-mitotic neurons under physiological conditions ^12,22,32^.

Using confocal microscopy, we reconstructed the morphology of 7 DIV control and REST-DBD neurons (Figure 4A). Accordingly, the expression of REST-DBD led to a significant reduction of axon and dendrite lengths (Figure 4A, C-E). Of note, the presence of varicosities along the axon was observed, unveiling the onset of a degenerative process (Figure 4A, black arrows). Thus, this data indicates that blocking the access to RE-1 sites in post-mitotic neurons leads to neuritic retraction.

**Figure 4.**
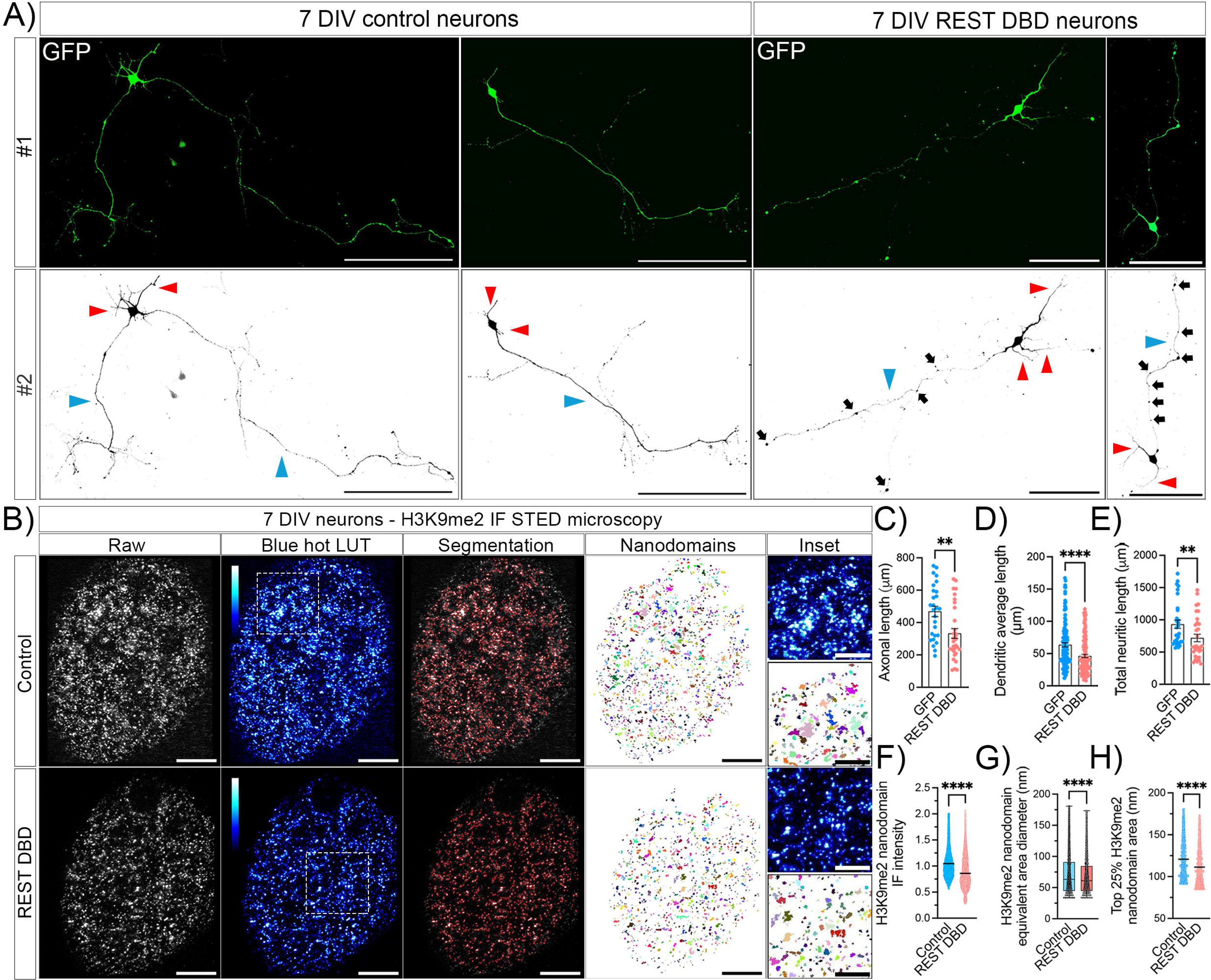
REST-DBD expression leads to neuritic retraction and dismantling of H3K9me2 nanodomains. **A)** Representative images of control and REST-DBD transfected neurons. Neurons were transfected 18 h after plating with control or REST- DBD construct, using GFP as a reporter. Then, they were fixed at 7 DIV and visualized by z-stack confocal imaging. **B)** Representative STED nanoscopy images showing H3K9me2 nuclear detection in control and REST-DBD neurons. **C, D, E)** Quantification of axonal (blue arrowheads), dendritic (red arrowheads) and total neuritic lengths in control and REST-DBD neurons. **F)** IF intensity measured in each H3K9me2 nanodomain of control and REST-DBD neurons. Mann-Whitney test, N=3. **G)** H3K9me2 equivalent area diameter of control and REST-DBD neurons (whole population). Kolmogorov-Smirnov test, N=3. **H)** Representation of the top 25% largest H3K9me2 nanodomains (equivalent area diameter of control and REST-DBD neurons). Kolmogorov-Smirnov test, N=3.

Additionally, both the intensity and topology of H3K9me2 nanodomains were analyzed (Figure 4B). The IF intensity was measured in each domain by STED microscopy, detecting a significant reduction in REST-DBD neurons. These observations were consistent with the confocal whole-nucleus z-stack imaging shown in Supplementary Figure 3C. Of note, the fold-change detected was higher using STED imaging than confocal microscopy, suggesting local changes at the nanoscale of heterochromatin.

The area of H3K9me2 nanodomains was also estimated (Figure 4B). Accordingly, the blockade of RE-1 sites dismantled H3K9me2 nanodomains, which was especially evident within the highest quartile of the population (top 25%, figure 4G, H). Statistical comparison between highest quartiles reported a significant shrinkage of H3K9me2 nanodomains in REST-DBD neurons (Figure 4H). These results suggest that RE-1 sites remain active in post-mitotic neurons, preserving H3K9me2 heterochromatin homeostasis and neuronal growth.

## Discussion

This article reports the contribution of CoREST-2 and RE-1 sites in maintaining H3K9me2 heterochromatin homeostasis and growth of neurons. On one hand, nuclear depletion of CoREST-2 leads to the enlargement of H3K9me2 nanodomains. On the other hand, the occupancy of RE-1 sites triggers the dissolution of H3K9me2 heterochromatin. In both cases, hippocampal neurons displayed neuritic retraction, affecting both axonal and dendritic compartments. These findings indicate that post-mitotic brain neurons demand a precise balance of H3K9me2 heterochromatin to preserve their growth and connectivity (Figure 5).

**Figure 5.**
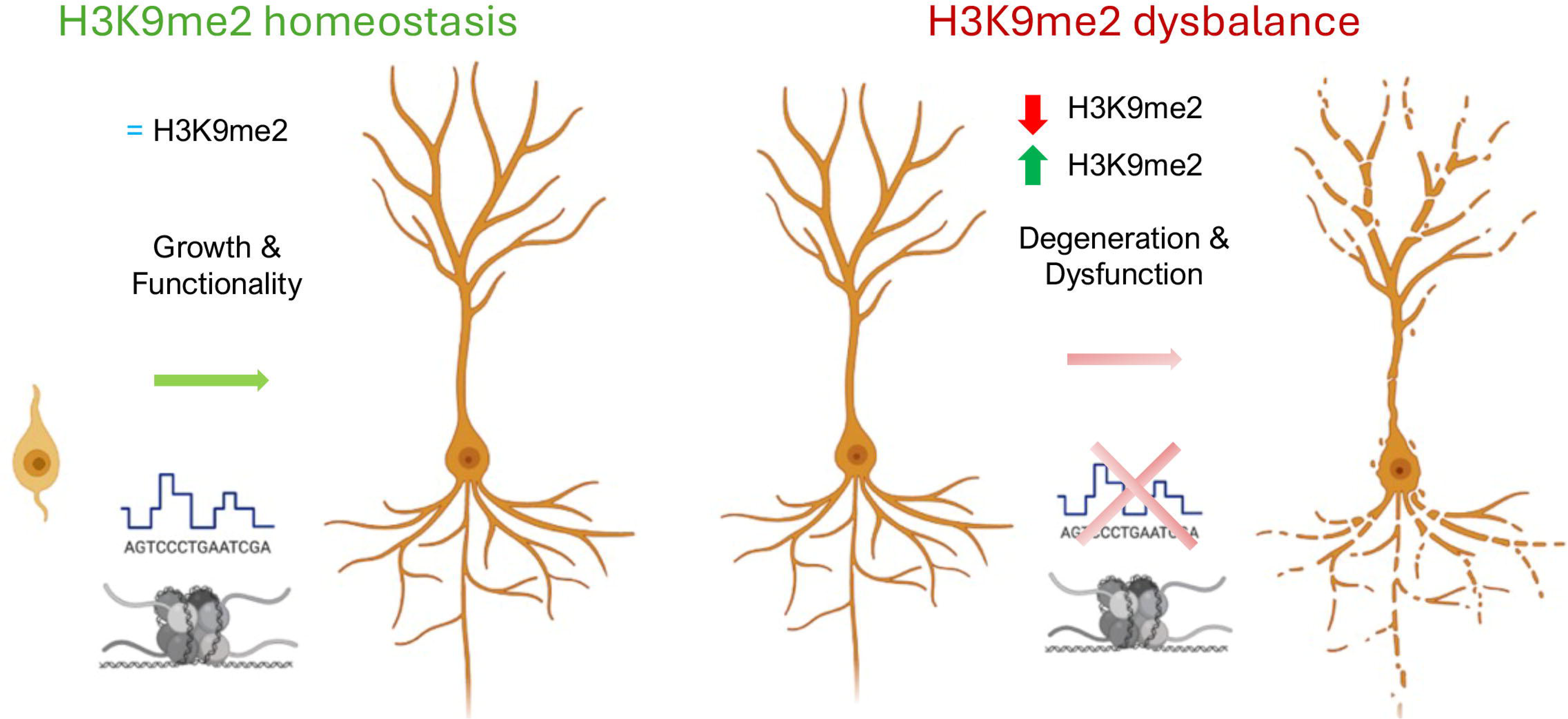
Proposed model for H3K9me2 heterochromatin homeostasis in growing neurons. The left side of the image illustrates a homeostatic situation, where H3K9me2 is balanced, and neurons grow properly. By contrast, the right side represents either up or down-regulation of H3K9me2 heterochromatin, which correlates with neuronal dysfunction characterized by a global reduction on neuritic growth.

The family of CoREST paralogues is involved in neuronal differentiation, although isoform-specific functions are just starting to emerge ^14,33^. For instance, CoREST-1 and - 2 are equally required to preserve the neurogenic niches of the brain cortex ^17,25^. Moreover, CoREST-1 is instrumental for the cortical migration of brain neurons ^15^. Nevertheless, the contribution of CoREST-2 to the development of post-mitotic neurons has remained understudied. In this work, we show that the genetic silencing of CoREST- 2 leads to a global neurite retraction, characterized by a reduction on axonal and dendritic lengths. Importantly, this phenotype suggests that endogenous CoREST-2 preserves the extension and plasticity of these compartments, which represents the structural basis for synapses and connectivity of brain neurons The CoREST paralogues are also linked to chromatin remodeling. Whilst CoREST-1 has been associated with H3K9 bi-methylation (H3K9me2) through G9a ^15,16,34,35^, CoREST-2 has been linked to H3K4me1/2 demethylation through LSD-1 ^17,36^. A short splicing variant of LSD1, namely LSD1+8a, demethylates H3K9me2 in the human neuroblastoma SH-SY5Y ^18^, but specific control of H3K9me2 by CoREST-2 in brain neurons had not been probed. In this regard, our data is instrumental to fill this gap, supporting that CoREST-2 promotes H3K9me2 demethylation and heterochromatin homeostasis in rat hippocampal neurons.

CoREST proteins do not have intrinsic enzymatic activity ^14^. Instead, they recruit factors catalyzing chemical modifications into chromatin. Therefore, probing its biochemical activity can be challenging. Biochemical analyses, such as immunoprecipitation followed by in vitro enzymatic assays, provide valuable insights but often lack of sub-cellular information. In this work, we visualized chromatin remodeling in cultured neurons *in situ*, using state-of-the-art microscopy setups, ranging from Airyscan 2 to STED imaging. Accordingly, we noted that H3K9me2 is organized in small clusters in the chromatin, here referred as nanodomains. Considering that H3K9me2 is a hallmark of heterochromatin, our results suggest that silencing CoREST-2 enlarges heterochromatin repressive domains and restrain neurite growth. Previous articles have shown that erasing H3K9me2, mostly by G9a inhibition, leads to neuritic retraction ^8,9^. Taken together, our findings support the notion that neurons demand a proper balance of H3K9me2 heterochromatin to preserve their growth and connectivity.

Intriguingly, a previous work reported a progressive decrease of CoREST-2 mRNA levels in 1-6 DIV rat neurons ^35^. By contrast, our data shows that CoREST-2 protein accumulates in the nuclei of neurons within a similar timeframe (1-7 DIV). However, both observations could be compatible since mRNAs and proteins may have divergent behaviors. Moreover, alternative splicing of CoREST-2 isoforms cannot be discarded within this timeframe, as reported for other H3K9me2 regulators such as G9a and LSD18,9,18.

In this work, we also tested the contribution of RE-1 sites to the post-mitotic life of neurons, focusing on growth and H3K9me2 heterochromatin. Epigenetic factors such as CoREST-1, MeCP2 and HDACs bind to RE-1 sites in the promoter of neuronal genes in fibroblast and stem cells ^22^. However, an *in-silico* analysis conducted in this work revealed that several non-neuronal genes carry RE-1 sites, covering global biological processes such as vesicle-mediated trafficking, cytoskeleton organization and cell death. These observations led us to hypothesize that RE-1 sites might remain active during post-mitotic stages of the neuron, contributing to chromatin remodeling and neuronal growth.

To block the access to RE-1 sites, neurons were transfected with the REST-DBD construct, according to previous works ^20,22,31^. Consequently, these neurons displayed axon and dendritic retraction, paralleled by the dismantling of H3K9me2 heterochromatin. Both the intensity and area of H3K9me2 nanodomains were significantly reduced after REST-DBD expression. These findings raised several hypotheses. For instance, additional epigenetic factors could bind to RE-1 sites. Along this line, G9a represses the expression of the RhoA-GEF Lfc in rat hippocampal neurons, a strong inhibitor of axon extension, by catalyzing H3K9me2 in its promoter region harboring a RE-1 site ^8^. Importantly, the neuronal phenotype obtained after suppressing G9a shares common features with the RE-1 blockade phenotype, such as axonal retraction and H3K9me2 erasure at chromatin ^8^. Moreover, previous reports described that RE-1 sites contain or are flanked by CpG islands, which makes them targets of DNA methylation ^16,22^. Seminal works in the field state that CpG methylation remains across different stages of neuronal development, working as spatial hints for chromatin repression ^16,22^. In this context, our results open a discussion about the role of RE-1 sites and their potential readers in post-mitotic neurons, as well as their contribution to controlling heterochromatin homeostasis and neuronal growth.

H3K9me2 is a hallmark of heterochromatin and when erased leads to chromatin relaxation, a conserved phenomenon in eukaryotic cells ranging from plants to animals ^37–40^. Heterochromatin is usually linked to transcriptional silencing, but its homeostasis contributes beyond gene expression. For instance, genomic stability depends on chromatin conformation. In particular, the accessibility of the DNA-repair machinery is tightly linked to H3K9me2-dependent heterochromatin ^41,42^. Therefore, unveiling mechanisms fine-tuning chromatin dynamics is at the base of neuronal physiology and disease. In this regard, the microscopy analyses addressed in this work enabled direct visualization of chromatin dynamics *in situ*. This approach could be highly informative for screening chromatin remodeling before conducting epigenomic and transcriptomic analyses.

In summary, this work proposes that CoREST-2 and RE-1 sites play a key role in fine-tuning H3K9me2 heterochromatin and supporting the growth of neurons. Chromatin works as a molecular memory, integrating information from the cellular and environmental context. This concept is particularly relevant for long-live cells like neurons, demanding robust strategies to preserve cognitive and motor functions of the nervous system.

## Materials and methods

### Reagents

Antibodies used in this work: anti-CoREST-2 (SIGMA-Merck, cat. n°: HPA021638), anti- H3K9me2 (Abcam, cat. n°: ab176882), anti-G9a (SIGMA, cat. n°: SAB2700645), anti-REST (Santa Cruz Biotechnologies, cat. n°: sc-374611), anti-MAP2 (BioLegend, cat. n°: 822501), anti-ß3-tubulin (Abcam, n° cat.: ab78078), anti-chicken Alexa Fluor 488 (ThermoFisher, cat. n°: a11039), anti-mouse Alexa Fluor 546 (ThermoFisher, cat. n°: a11010), anti-rabbit Alexa Fluor 568 (ThermoFisher, cat. n°: a11004), anti-rabbit Alexa Fluor 633 (ThermoFisher, cat. n°: a21070), anti-rabbit Fab Alexa Fluor 594 (Jackson Laboratories, cat. n°: 111-587-003), anti-rabbit Fab Alexa Fluor 647 (Jackson Laboratories cat. n°: 111-607-003), anti-rabbit Atto 633 (SIGMA, cat. n°: 77671). Neurobasal, B27, Glutamax, Pen/Strep, Optimem, and Lipofectamine 2000 reagent were purchased to ThermoFisher (USA). Plasmid DNA purifications were done using commercial kits (Qiagen). All other reagents used in this work were analytical grade.

### Culture of embryonic hippocampal rat neurons

Hippocampal neurons isolated from embryonic (E18,5) rat brains were isolated and cultured according to established protocols ^43,44^. Briefly, a pregnant E18,5 Wistar rat was euthanized and embryos removed for brain isolation. Hippocampi were dissected and collected into phosphate buffer saline (PBS), followed by enzymatic digestion (trypsin-PBS solution, 20 min at 37°C). Then, tissue was mechanically disaggregated until no clumps were observed (by up-and-down using p1000 and p200 micropipettes). Then, cells were counted using a Neubauer cell chamber to estimate the yield. For IF experiments, 6×10^4^ neurons were plated on 12 mm diameter glass coverslips, pre-treated with poly-L-lysine, using 24-well plates. Neurons were plated for 1h in MEM supplemented with 10% horse serum. Then, media was discarded and replaced by Neurobasal medium supplemented with B27, Glutamax and penicillin + streptomycin. This media was refilled every 3 days to preserve osmolarity, maintaining 2/3 of conditioned media volume. Rats were purchased at the animal facility of the Instituto Ferreyra (INIMEC-CONICET, UNC – Cordoba, Argentina) and handled following local and international guidelines for the use of laboratory animals.

### ImmunoWluorescence (IF) for intracellular protein detection

Neurons were cultured for the times indicated in each figure and fixed using 4% paraformaldehyde/sucrose-PBS solution (20 min, room temperature, RT). Then, neurons were washed 3 times with PBS (5 min each, RT), permeabilized using 0.2% triton x-100/PBS for 10 min at RT and blocked with 1% BSA dissolved in 0,1% Tween- 20/PBS (PBST) for 1h at RT. Primary antibodies were diluted in 0.1% PBST solution and incubated ON at 4°C inside a wet-air chamber to avoid evaporation (16-18 h). Coverslips were washed 3 times with 0.1% PBST (5 min each) and Alexa Fluor (AF)-conjugated secondary antibodies were incubated for 1h at RT (1:1.000, 0.1% PBST). Consequently, coverslips were washed 1 time with 0.1% PBST, followed by DAPI-PBS incubation (5 min, RT) and final 5 min wash using PBS (RT). Coverslips-containing neurons were mounted in glass slides using Mowiol. Once dry, slides were stored at 4°C in darkness until imaging. Of note, IF staining for 1-14 DIV time-course assays were made only after collecting all times to avoid artifacts attributable to unintentional aleatory errors on pipetting or antibody dilution (Figures 1A-D, 2A-F, 5A-H). Accordingly, fixed coverslips- containing neurons were maintained at 4°C under sterile conditions until completing time courses.

### Nuclear and cytosolic detection of proteins

IF samples were imaged using a LSM 800 Zeiss confocal microscope, under z-stack configuration (63x oil objective 1.4 NA, 1.2 zoom in; pinhole: 1 airy unit (AU); 1 µm step size). Samples were excited using 405, 488, and 546 nm wavelength laser according to each fluorochrome. For time-course analysis (1-14 DIV neurons), the following combination of antibodies and fluorochromes were used: DAPI (405 nm); anti-MAP2 (chicken) + anti-chicken AF 488; CoREST2/G9a/H3K9me2 (rabbit) + anti-rabbit AF 546. Then, z-stack images were reconstructed by post-imaging analysis in Fiji-ImageJ, using the sum slices z-project, followed by background subtraction. The nuclear region was determined by the DAPI channel, drawing a region of interest (ROI) and then estimating raw integrated density (ID). Using the MAP2 channel, a ROI was drawn in the whole soma (without neurites) and ID was calculated. Accordingly, cytosolic IF signal resulted from subtracting the ‘whole soma ROI’ minus ‘nuclear soma ROI’. Then, the ratio ‘nucleus/cytosol’ was determined as a measurement of nuclear enrichment.

### Molecular cloning of shRNA CoREST-2

The small hairpin RNA (shRNA) targeting rat and human CoREST-2 mRNA was built using the pB6/U6 plasmid in two steps ^8^. The sequence chosen was GGGGATGCTGGTGTGGTCACC. For cloning, the following 5’ to 3’ oligonucleotides were synthetized: 1a: GGGATGCTGGTGTGGTCACCA; 1b: AGCTTGGTGACCACACCAGCATCCC; 2a: AGCTTGGTGACCACACCAGCATCCCCTTTTTG; 2b: AATTCAAAAAGGGGATGCTGGTGTGGTCACCA. The plasmid pB6U6 was enzymatically digested by Apa, Klenow and HindIII (AKH) and purified using the GeneJET gel extraction kit (Thermo Fisher). Oligonucleotides were resuspended in DNA/RNAse free milli-Q water. The pair 1a+1b was annealed and then ligated to pB6U6-AKH for 1h at RT. DH5a bacteria were transformed using 2 µL of ligation product (ON, 37°C). Next day, 8- 10 colonies were picked for replica plates and colony PCR screening (Fw primer: T7; Rv primer: oligo 1b). Positive colonies were cultured ON, harvested and plasmid DNA purified by minipreps (pB6U6 1^st^). Then, pB6U61^st^ was digested by EcoRI and HindIII (E/H) and purified accordingly. The oligonucleotide pair 2a+2b was annealed and ligated to the pB6U6 1^st^-E/H. DH5a bacteria were transformed using 2 µL of ligation product. Colonies were picked and cultured ON for minipreps. Plasmids were then sequenced in Macrogen (Korea) to check correct shRNA assembly. Finally, the shRNA cassette was released using SpeI restriction enzyme and subcloned into the pCAGIG vector, encoding GFP as a fluorescent reporter.

### Transient transfection of rat neurons

Neurons were transfected 18 h after plating with Lipofectamine 2000 and Opti-MEM. Accordingly, neuronal culture media (Neurobasal + supplements) was removed and replaced by 200 µL Opti-MEM per well (24 multi-well). Neuronal culture media was stored at 4°C under sterile conditions until completing transfection. For one well reaction, 500 ng of DNA were resuspended in 25 µL Opti-MEM (Tube A), whereas 1 µL Lipofectamine was resuspended in an additional 25 µL Opti-MEM (Tube B). Scaling-up was done following this ratio. After 5 min, Tube A and Tube B were mixed and gently shake every 5 min until completing 15 min. Then, 50 µL of transfection mixture (DNA + Lipofectamine) were added per well, previously containing 200 µL Opti-MEM. For CoREST-2 KD experiments, 500 ng of shRNA sc or shRNA CoREST-2 plasmids were used per well. For REST DBD assays, 100 ng of pCAGIG (GFP reporter) + 400 ng empty pcDNA or 400 ng REST-DBD construct were used. After 3 h, the reaction was stopped by discarding the transfection mixture and restoring neuronal culture media collected before transfection. Transfected neurons were cultured until completing 7 DIV, refilling media every 3 days.

### Morphometric analysis of neurons

For neuritic length measurements, 7 DIV GFP-transfected neurons (control, CoREST KD or REST-DBD) were imaged using a LSM 800 Zeiss confocal microscope, under z-stack configuration (20x objective, 0.8 NA; pinhole: 1 AU; 1 µm step size). GFP-expressing neurons were completely reconstructed by imaging several fields, which were stitched post-imaging using the ‘stitching’ plug-in of Fiji (‘*pairwise stitching’*). Maximal intensity was projected in each case to improve GFP signal, and background subtracted. Images shown in Figure 1G and 4A correspond to GFP and grayscale binary masks, used for neuritic length measurements. In some cases, GFP-expressing neurons were impossible to reconstruct because low signal, especially towards the medial-to-distal axon. Thus, neurons were discarded from the analysis. Axonal domain is clearly distinguished because its length is at least 2-3 times of dendrites, according to established protocols. Dendritic average length corresponds to the mean length of all dendrites per neuron. Total neuritic length reflects the sum of all neurites (axon + dendrites).

### Colocalization assays

To analyze co-distribution between CoREST-2, H3K9me2 and G9a, 7 DIV rat hippocampal neurons were used. They were stained as indicated above, but monovalent secondary antibodies conjugated to Alexa Fluor probes were used (Fab fragments, Jackson Laboratories, dilution 1:200). Coverslips-containing neurons were ON incubated using anti-MAP2 and anti-CoREST2. Then, they were washed 3 times using 0.1% PBST. Secondary antibodies anti-chicken AF 488 and anti-rabbit Fab-AF 546 were incubated 1h at RT (0.1% PBST). Then, coverslips were washed 3 times with 0.1% PBST and post-fixed using 4% PFA/sucrose-PBS solution for 10 min, followed by 0.1 M glycine for 10 min at RT. Then, either H3K9me2 or G9 antibodies were incubated for 1h at RT, followed by 3 washes with 0.1% PBST. Accordingly, secondary anti-rabbit Fab-AF 633 was incubated for 1 h in 0.1% PBST. Finally, coverslips were washed 1 time with 0.1% PBST, incubated with DAPI-PBS solution for 5 min and finally washed with PBS (5 min). Coverslips were mounted in glass slides using Mowiol.

Imaging was performed using a LSM 980 AiryScan 2 confocal microscope, under z-stack and super resolution (SR) configuration, improving by 1.4x spatial resolution compared to conventional confocal microscopy (objective: 63x oil, 1.4 NA; pinhole: 1 AU; step size: 120 nm). After imaging, maximal intensity was projected using z-project of Fiji. Background was subtracted and median filter was applied. Images were deconvoluted using the ‘3D deconvolution lab’ plug-in of Fiji. Pearson’s and Mander’s coefficients were estimated by the ‘coloc-2’ plug-in in nuclear regions (delimited by DAPI), using an automatized macro-based pipeline. The meaning of M1 and M2 Mander’s coefficients are explained in Figure 2I, J.

### Characterization of H3K9me2 nanodomains by STED microscopy

Seven DIV control, CoREST-2 KD, and REST-DBD neurons were imaged using a STEDYCON STED microscope (Abberior Instruments) equipped with a 100× objective (NA = 1.4). Immunofluorescence staining was performed as described in the Methods section, with modifications including a higher concentration of the primary anti- H3K9me2 antibody (1:200) and the secondary anti-rabbit Atto594-conjugated antibody at 1:500. DAPI staining was omitted. Images were acquired at a pixel size of 15 nm, with each line scanned 10 to 20 times (line accumulations). Excitation was achieved using a 561-nm laser, and depletion was performed with a 775-nm laser.

Automated image analysis was conducted using a custom pipeline developed in CellProfiler1 (www.cellprofiler.org) to quantify the area and intensity of H3K9me2 nanodomains. STED images were batch processed. Briefly, threshold was adjusted in each image using an adaptive strategy based on the “robust background” method, followed by watershed segmentation to create a mask with individual nanodomains. Domains smaller than four pixels were excluded from the analysis. The remaining domains were filled to correct for segmentation defects. Pixel intensity for each domain was then measured in the original images ^45^.

### IdentiWication of Genes Harboring RE-1 sites and Functional Annotation

The motifs matrices MA0138.1, MA0138.2, and MA0138.3, representing RE-1 binding sites, were retrieved from the JASPAR database and used for sequence analysis ^46^. Binding site prediction was performed using the FIMO tool from the MEME suite (version 5.5.7) against the rn6 rat genome assembly ^47^. While scanning for MA0138.1 and MA0138.3 did not yield significant results, analysis of MA0138.2 identified 75,725 potential transcription factor binding sites (TFBS). The resulting TFBS were intersected with genomic annotations from the RefSeq database, refining the dataset to 26,014 TFBS associated with gene promoters. Promoter regions were defined as sequences within 1,500 base pairs (bp) upstream of transcription start sites (TSS). This filtering step identified 8,776 genes with potential RE.1 binding sites in their promoters. Further selection was performed using statistical significance thresholds, retaining genes with a q-value < 0.05, resulting in a final set of 637 genes. Gene Ontology (GO) analysis was conducted to classify these genes into Biological Process (BP) and Molecular Function (MF) categories, enabling the characterization of their functional roles.

### Gene Ontology (GO) Annotation and Visualization

Gene Ontology (GO) analysis was conducted for the final set of 637 genes using the GOTermMapper tool (https://go.princeton.edu/cgi-bin/GOTermMapper) to classify them into Biological Process (BP) and Molecular Function (MF) categories ^48^. GO terms associated with more than 50 genes for BP and more than 10 genes for MF were selected for downstream analysis. The selected GO terms were visualized as treemaps using the treemap package (version 2.4-4) in R. Each treemap represents GO terms as hierarchical rectangles, where the size of each rectangle is proportional to the number of genes annotated to the corresponding term. A color gradient was applied to indicate the gene counts, providing a clear visualization of the most enriched GO categories.

## Statistical analysis

Statistics were calculated in GraphPad Prism 10. Gaussian distribution was determined by the Shapiro-wilk normality test. Tests and post-tests are indicated in figure legends. Plots represent the mean ± standard error median (SEM), excepting box plots of Figures 3C and 4G. Outliers were cleaned from raw data using a Rout test (0.1%). Experimental points considered for the analysis are shown in each plot. Results represent the trend of at least 3 independent cultures (N=3). Statistical significance was set for p-values below 0.05 (*<0.05, **<0.01, ***< 0.001, ****0.0001).

## Figure rendering

Figures were drawn using Adobe Photoshop. Model proposed in Figure 5 was drawn in Biorender and Microsoft Office Power Point.

## Supporting information

https://drive.google.com/drive/folders/1ja9a01SJsawPYnmCOjCj_GlckqccLCZL?usp=drive_link

## Acknowledgments

This work was funded by the National Council for Scientific Research of Argentina (CONICET, PIP n° to AC, ALM, AC-G and CW), the National Agency for Science (ANPCyT, PICT 2019-00471, PICT 2021-00630 to CW) and the International Society for Neurochemistry (CAEN 1B to CW). AC-G, VR-S, MR, DC, ALM, AC and CW are staff scientists of CONICET. JFD, IG and LS were students of Biology at the National University of Córdoba (Argentina) by the time of this study. We especially thank Dr. Gail Mandel for providing us with the REST-DBD construct, and Dr. Michael Spinner for kindly delivering it (Oregon & Health Science University, USA). We also acknowledge Dr. Cecilia Sampedro, Dr. Gonzalo Quasollo, and Dr. Carlos Mas for technical assistance at the Center for Microscopy and Nanoscopy of Córdoba (CEMINCO, National University of Córdoba). We thank Silvina Ferrer (INIMEC) and Candy Bravo (FUCIBICO) for grant administration.

---

**Supplementary Figure 1.**
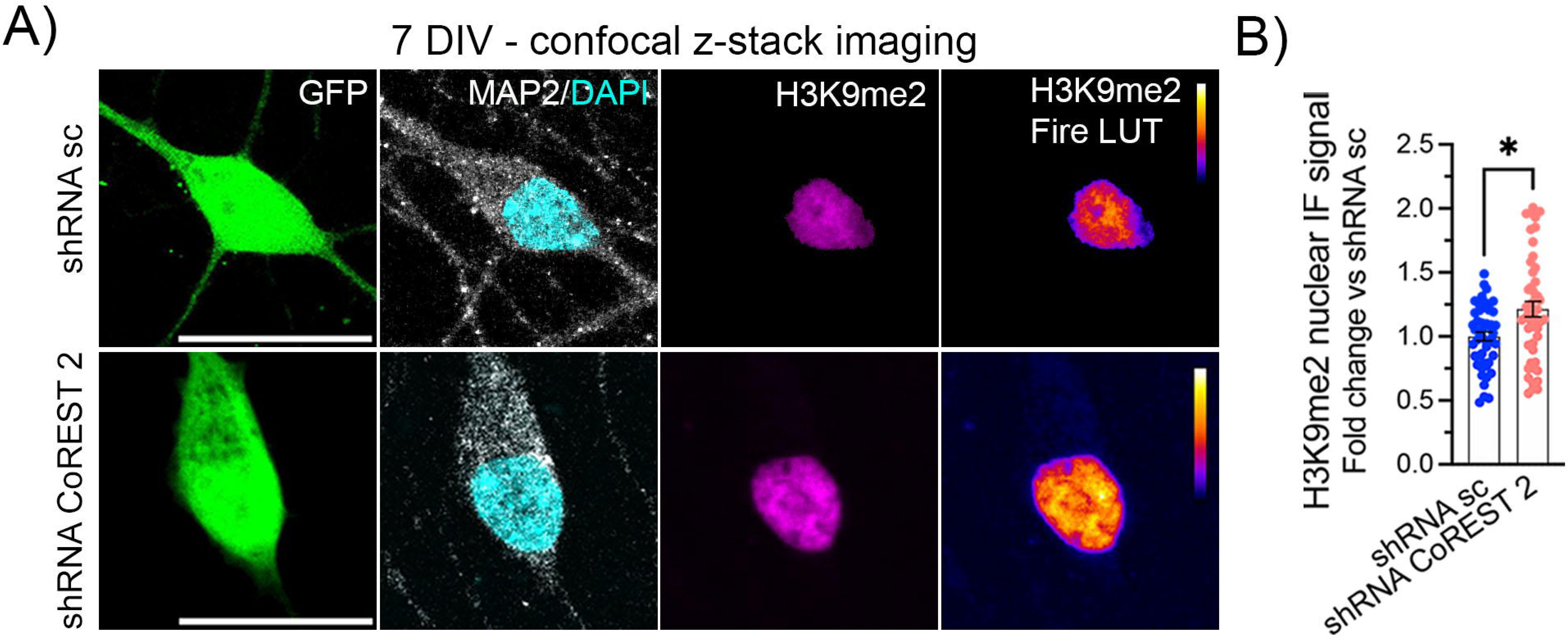
**Confocal z-stack maximal intensity of H3K9me2 in control and CoREST-2 KD neurons**. Representative image **(A)** and quantification **(B)** of H3K9me2 using z-stack confocal imaging to capture whole nuclear signal. Neurons were transfected 18 h after plating, fixed at 7 DIV, and stained to detect H3K9me2. Mann-Whitney test, N=3. Scale: 10 µm.

**Supplementary Wigure 2.**
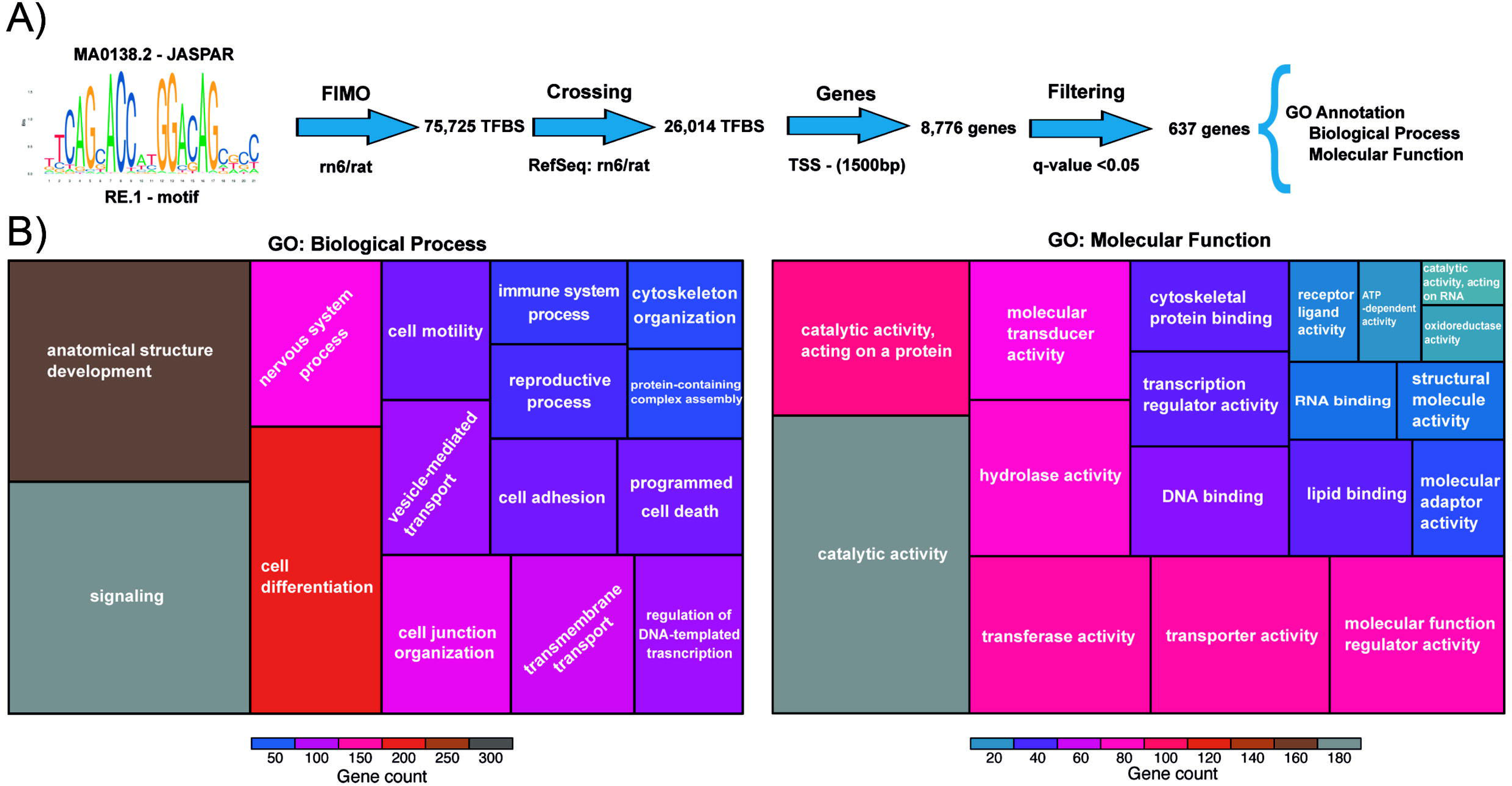
Gene ontology of RE-1 sites. **A)** Pipeline for RE-1 sites analysis using FIMO tool. **B)** Treemaps illustrating Gene Ontology (GO) terms for the Biological Process (BP) and Molecular Function (MF) categories. Terms associated with more than 50 genes in BP and more than 10 genes in MF are shown for each category.

**Supplementary Figure 3.**
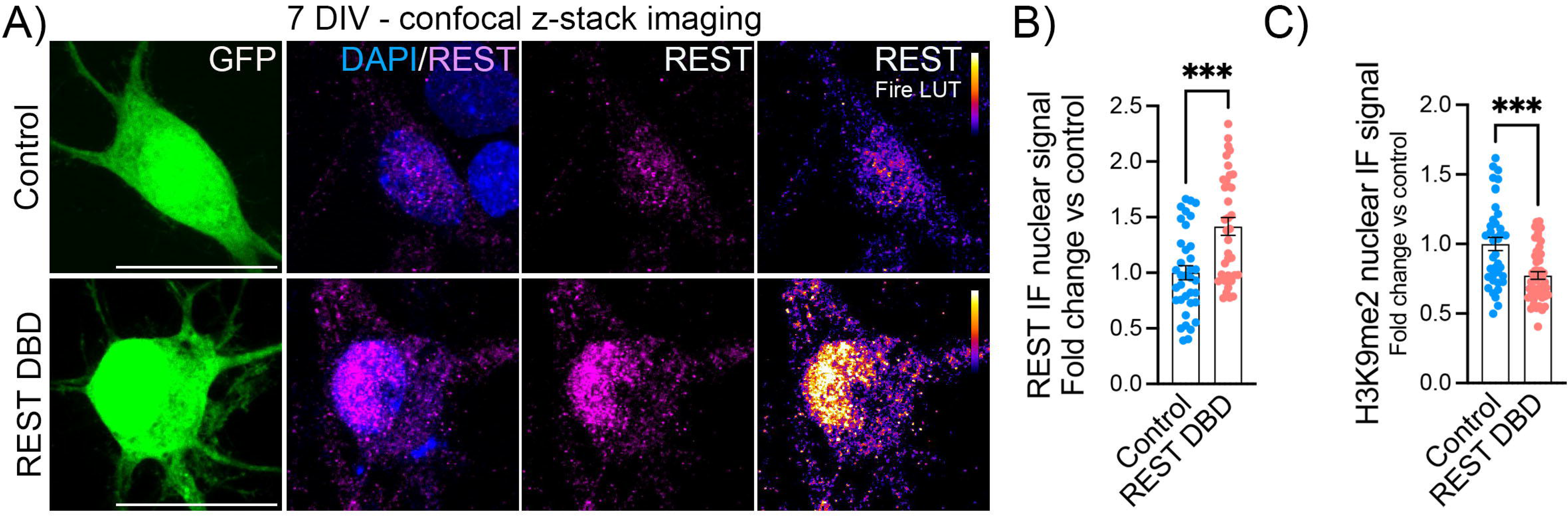
**Control of expression of REST-DBD construct and H3K9me2 confocal microscopy quantiWication**. **A)** Representative confocal z-stack maximal intensity of control and REST-DBD expressing neurons. Cells were transfected 18h after plating and fixed at 7 DIV, followed by IF staining and confocal imaging. Scale: 10 µm. **B)** Quantification of total REST IF nuclear signal in control and REST-DBD neurons. Mann-Whitney test, N=3. **C)** Quantification of whole-nucleus z-stack imaging of H3K9me2 IF signal in control and REST-DBD neurons. Neurons were transfected 18 h after plating, fixed at 7 DIV, and stained to detect H3K9me2. Mann-Whitney test, N=3.

## Notes

### Competing Interest Statement

The authors have declared no competing interest.

## References

1. Abay-Nørgaard, S., Attianese, B., Boreggio, L. & Salcini, A. E. Regulators of H3K4 methylation mutated in neurodevelopmental disorders control axon guidance in Caenorhabditis elegans. Development 147, (2020).

2. Mariani, L., Lussi, Y. C., Vandamme, J., Riveiro, A. & Salcini, A. E. The H3K4me3/2 histone demethylase RBR-2 controls axon guidance by repressing the actin-remodeling gene wsp-1. Development (Cambridge) 143, 851–863 (2016).

3. Riveiro, A. R. et al. JMJD-1.2/PHF8 controls axon guidance by regulating hedgehoglike signaling. Development (Cambridge) 144, 856–865 (2017).

4. Wilson, C. & Cáceres, A. New insights on epigenetic mechanisms supporting axonal development: histone marks and miRNAs. FEBS Journal 288, 6353–6364 (2021).

5. Lindner, R., Puttagunta, R. & Di Giovanni, S. Epigenetic Regulation of Axon Outgrowth and Regeneration in CNS Injury: The First Steps Forward. Neurotherapeutics 10, 771–781 (2013).

6. Gaub, P. et al. HDAC inhibition promotes neuronal outgrowth and counteracts growth cone collapse through CBP/p300 and P/CAF-dependent p53 acetylation. Cell Death Differ 17, 1392–1408 (2010).

7. Wilson, C., Moyano, A. L. & Cáceres, A. Perspectives on Mechanisms Supporting Neuronal Polarity From Small Animals to Humans. Frontiers in Cell and Developmental Biology vol. 10 Preprint at 10.3389/fcell.2022.878142 (2022).

8. Wilson, C. et al. The Histone Methyltransferase G9a Controls Axon Growth by Targeting the RhoA Signaling Pathway. Cell Rep 31, (2020).

9. Fiszbein, A. et al. Alternative Splicing of G9a Regulates Neuronal Differentiation. Cell Rep 14, 2797–2808 (2016).

10. Roopra, A., Qazi, R., Schoenike, B., Daley, T. J. & Morrison, J. F. Localized domains of G9a-mediated histone methylation are required for silencing of neuronal genes. Mol Cell 14, 727–738 (2004).

11. Tachibana, M., Matsumura, Y., Fukuda, M., Kimura, H. & Shinkai, Y. G9a/GLP complexes independently mediate H3K9 and DNA methylation to silence transcription. EMBO Journal 27, 2681–2690 (2008).

12. Hwang, J. Y., Aromolaran, K. A. & Zukin, R. S. The emerging field of epigenetics in neurodegeneration and neuroprotection. Nat Rev Neurosci 18, 347–361 (2017).

13. María, M., et al. CoREST: A Functional Corepressor Required for Regulation of Neural-SpeciKic Gene Expression. Neurobiology Communicated by William J. Lennarz vol. 96 www.pnas.org. (1999).

14. Maksour, S., Ooi, L. & Dottori, M. More than a corepressor: The role of corest proteins in neurodevelopment. eNeuro vol. 7 Preprint at 10.1523/ENEURO.0337-19.2020</year> (2020).

15. Fuentes, P., Cánovas, J., Berndt, F. A., Noctor, S. C. & Kukuljan, M. CoREST/LSD1 control the development of pyramidal cortical neurons. Cerebral Cortex 22, 1431–1441 (2012).

16. Ballas, N. & Mandel, G. The many faces of REST oversee epigenetic programming of neuronal genes. Current Opinion in Neurobiology vol. 15 500–506 Preprint at 10.1016/j.conb.2005.08.015 (2005).

17. Wang, Y. et al. LSD1 co-repressor Rcor2 orchestrates neurogenesis in the developing mouse brain. Nat Commun 7, (2016).

18. Laurent, B. et al. A Specific LSD1/KDM1A Isoform Regulates Neuronal Differentiation through H3K9 Demethylation. Mol Cell 57, 957–970 (2015).

19. Lunyak, V. V. & Rosenfeld, M. G. No rest for REST: REST/NRSF regulation of neurogenesis. Cell 121, 499–501 (2005).

20. Chong, J. A., et al. REST: A Mammalian Silencer Protein That Restricts Sodium Channel Gene Expression to Neurons. Cell vol. 80 (1995).

21. Schoenherr, C. J., Paquette, A. J. & Anderson, D. J. Identification of potential target genes for the neuron-restrictive silencer factor. Proc Natl Acad Sci U S A 93, 9881–9886 (1996).

22. Ballas, N., Grunseich, C., Lu, D. D., Speh, J. C. & Mandel, G. REST and its corepressors mediate plasticity of neuronal gene chromatin throughout neurogenesis. Cell 121, 645–657 (2005).

23. 23. Li, L., Suzuki, T., Morit, N. & Greengard, P. *IdentiKication of a Functional Silencer Element Involved in Neuron-SpeciKic Expression of the Synapsin I Gene*. *Proc*. Natl. Acad. Sci. USA vol. 90 https://www.pnas.org (1993).

24. 24. Bruce, A. W., et al. Genome-Wide Analysis of Repressor Element 1 Silencing Transcription Factorneuron-Restrictive Silencing Factor (RESTNRSF) Target Genes. vol. 101 www.hgmp.mrc.ac.ukSoftwareEMBOSS (2004).

25. Monaghan, C. E. et al. REST corepressors RCOR1 and RCOR2 and the repressor INSM1 regulate the proliferation- differentiation balance in the developing brain. Proc Natl Acad Sci U S A 114, E406–E415 (2017).

26. Rozés-Salvador, V., Wilson, C., Olmos, C., Gonzalez-Billault, C. & Conde, C. Fine- Tuning the TGFβ Signaling Pathway by SARA During Neuronal Development. Front Cell Dev Biol 8, 1–13 (2020).

27. Rivera, C. et al. Revealing RCOR2 as a regulatory component of nuclear speckles. Epigenetics Chromatin 14, (2021).

28. Jacquemet, G., Carisey, A. F., Hamidi, H., Henriques, R. & Leterrier, C. The cell biologist’s guide to super-resolution microscopy. J Cell Sci 133, (2020).

29. Willig, K. I. et al. Nanoscale resolution in GFP-based microscopy. Nat Methods 3, 721–723 (2006).

30. Lunyak, V. V et al. Corepressor-Dependent Silencing of Chromosomal Regions Encoding Neuronal Genes. www.sciencemag.org.

31. Ballas, N. et al. Regulation of neuronal traits by a novel transcriptional complex. Neuron 31, 353–365 (2001).

32. McGann, J. C. et al. The genome-wide binding profile for human RE1 silencing transcription factor unveils a unique genetic circuitry in hippocampus. Journal of Neuroscience 41, 6582–6595 (2021).

33. Maksour, S. et al. REST and RCOR genes display distinct expression profiles in neurons and astrocytes using 2D and 3D human pluripotent stem cell models. Heliyon 10, (2024).

34. Ceballos-Chávez, M., et al. Control of neuronal differentiation by sumoylation of BRAF35, a subunit of the LSD1-CoREST histone demethylase complex. Proc Natl Acad Sci U S A 109, 8085–8090 (2012).

35. Sáez, J. E., et al. Decreased expression of CoREST1 and CoREST2 together with LSD1 and HDAC1/2 during neuronal differentiation. PLoS One 10, (2015).

36. Shi, Y. et al. Histone demethylation mediated by the nuclear amine oxidase homolog LSD1. Cell 119, 941–953 (2004).

37. 37. Yabe, K., et al. H3K9 Methylation Regulates Heterochromatin Silencing through Incoherent Feedforward Loops. Sci. Adv vol. 10 https://www.science.org (2024).

38. Montavon, T. et al. Complete loss of H3K9 methylation dissolves mouse heterochromatin organization. Nat Commun 12, (2021).

39. Fukuda, K. et al. Regulation of mammalian 3D genome organization and histone H3K9 dimethylation by H3K9 methyltransferases. Commun Biol 4, (2021).

40. Methot, S. P. et al. H3K9me selectively blocks transcription factor activity and ensures differentiated tissue integrity. Nat Cell Biol 23, 1163–1175 (2021).

41. Ciccia, A. & Elledge, S. J. The DNA Damage Response: Making It Safe to Play with Knives. Molecular Cell vol. 40 179–204 Preprint at 10.1016/j.molcel.2010.09.019 (2010).

42. Sun, Y. et al. Histone H3 methylation links DNA damage detection to activation of the tumour suppressor Tip60. Nat Cell Biol 11, 1376–1382 (2009).

43. Kaech, S. & Banker, G. Culturing hippocampal neurons. Nat Protoc 1, 2406–2415 (2006).

44. Wilson, C., Rozés-Salvador, V. & Cáceres, A. Protocol for Evaluating Neuronal Polarity in Murine Models. STAR Protoc 1, 100114 (2020).

45. Stirling, D. R. et al. CellProfiler 4: improvements in speed, utility and usability. BMC Bioinformatics 22, (2021).

46. Wasserman, W. W. & Sandelin, A. Applied bioinformatics for the identification of regulatory elements. Nature Reviews Genetics vol. 5 276–287 Preprint at 10.1038/nrg1315 (2004).

47. Grant, C. E., Bailey, T. L. & Noble, W. S. FIMO: Scanning for occurrences of a given motif. Bioinformatics 27, 1017–1018 (2011).

48. Boyle, E. I. et al. GO::TermFinder - Open source software for accessing Gene Ontology information and finding significantly enriched Gene Ontology terms associated with a list of genes. Bioinformatics 20, 3710–3715 (2004).

